# Multi-omics consensus ensemble refines the classification of muscle-invasive bladder cancer with stratified prognosis, tumour microenvironment and distinct sensitivity to frontline therapies

**DOI:** 10.1101/2021.05.30.446369

**Authors:** Xiaofan Lu, Jialin Meng, Liwen Su, Liyun Jiang, Haitao Wang, Junkai Zhu, Mengjia Huang, Wenxuan Cheng, Li Xu, Xinjia Ruan, Yujie Zhou, Shuyuan Yeh, Chaozhao Liang, Fangrong Yan

## Abstract

The molecular classification of muscle-invasive bladder cancer (MIBC) based on transcriptomic signatures has been extensively studied. The complementary nature of information provided by different molecular profiles motivated us to refine MIBC classification by aggregating multi-omics data. We generated a consensus ensemble through ten multi-omics integrative clustering approaches on 396 MIBCs from TCGA. A total of 701 MIBCs from different sequencing technologies were used for external validation. Associations between subtypes and prognosis, molecular profiles, the tumour microenvironment, and potential response to frontline therapies were further analyzed. Nearest template prediction and random forest classification were used to develop a predictive signature/classifier for MIBC refinement. We identified four integrative consensus subtypes of MIBC, which were further designated basal-inflamed, basal-noninflamed, luminal-excluded and luminal-desert by immune profiling. Of note, the refinement of basal-like MIBC classification adds to the literature by identifying a basal-noninflamed MIBC subtype presenting with a significantly poor outcome and a global immune-cold phenotype, which might be triggered by Chr4 deletion and high activation of the oncogenic *NRF2* pathway. In contrast, basal-inflamed MIBC showed high immunocyte infiltration and high expression of potential targets for immunotherapy. Using an external metastatic MIBC cohort in which patients received anti-PD-L1 treatment, we suggested that basal-inflamed MIBC had a higher likelihood of responding to immunotherapy than other MIBCs. The R package “*refineMIBC*” was offered as a research tool to refine MIBC from a single-sample perspective (https://github.com/xlucpu/refineMIBC). This consensus ensemble refines the intrinsic MIBC subtypes, which provides a blueprint for the clinical development of rational targeted and immunotherapeutic strategies.

## INTRODUCTION

Bladder cancer (BCa), the ninth most prevalent cancer, led to approximately 549,233 new cases and 199,922 related deaths globally in 2018 [1]. Approximately 75% of BCa patients are diagnosed with non-muscle-invasive bladder cancer (NMIBC) and are treated with transurethral resections accompanied by intravesical therapy with Bacillus Calmette-Guérin or chemotherapeutic agents. However, NMIBC reoccurs within 10 years in nearly three-fourths of patients with high-grade NMIBC, and 33% of these recurrent tumours progress to the muscle-invasive type (MIBC). In patients with MIBC, the 5-year overall survival (OS) rate is as low as 62-68% after radical cystectomy or chemotherapy and further decreases to 35% if metastasis to nearby lymph nodes or distant organs occurs.

MIBC is a heterogeneous disease characterized by genomic instability and a high mutation burden. Molecular profiling facilitates the classification of MIBC into diverse molecular subtypes, contributing to more precise guidance for prognosis prediction and therapeutic options. Several kinds of molecular classification of MIBC have been proposed based on various gene expression signatures at the transcriptome level [2-6], which has remarkably advanced the understanding of MIBC biology. However, none of the single molecular level approaches thoroughly captures the complicacy of the cancer genome or precisely pinpoints the oncogenic-driving mechanism, thus leading to an evolving scheme for classifying MIBC based on multi-omics profiles, which could have the latent to unveil further biological insights. Recently, Mo *et al*. reported for the first time the multi-omics classification of MIBC using a fully Bayesian latent variable integrative model and revealed the crosstalk between different molecular characteristics, resulting in only two subtypes (*i*.*e*., basal and luminal/differentiated), which may hardly decipher the heterogeneity of MIBC [7].

The Cancer Genome Atlas (TCGA) has made large-scale efforts to assemble transcriptomic, genomic and epigenomic profiles for over 400 MIBC cases, which offers a major opportunity to comprehensively delving MIBC subtypes. Given that more than a dozen multi-omics integration strategies have been proposed and applied in medical research [8], we aimed to identify integrative consensus subtypes (iCS) of MIBC by generating a consensus ensemble from the classifications obtained with different algorithms using transcriptomic protein-coding mRNA and long non-coding RNA (lncRNA) expression, genomic mutation and copy number alteration (CNA), and epigenomic DNA methylation profiles to provide better overview of tumour heterogeneity and biological processes.

## MATERIALS AND METHODS

### Multi-omics data sets

Molecular profiles of the TCGA-BLCA dataset were retrieved as MIBC-TCGA cohort used for the multi-omics data analysis [5], including 396 primary muscle-invasive bladder cancers with complete transcriptome expression, somatic mutations, CNAs, DNA methylation and survival outcomes available. For raw count data of high-throughput sequencing downloaded by the R package “*TCGAbiolinks*” [9], both mRNAs and lncRNAs were considered. Specifically, lncRNAs were identified according to Vega (http://vega.archive.ensembl.org/) for the following types: non_coding, 3prime_overlapping_ncRNA, antisense_RNA, lincRNA, sense_intronic, sense_overlapping, macro_lncRNA, and bidirectional_promoter_lncRNA. Ensembl IDs for transcriptomes were transformed into gene symbols by GENCODE27 mapping. The number of fragments per kilobase million (FPKM) was computed and converted into transcripts per kilobase million (TPM), which showed more similarity to the numbers obtained from microarray analysis and improved comparability between samples. DNA methylation profile was downloaded from the XENA database (https://xenabrowser.net/). Copy number segment data were collected from FireBrowse (http://firebrowse.org/). Somatic mutations, clinicopathological features, overall survival (OS) and progression-free survival (PFS) rate data were downloaded from cBioPortal (https://www.cbioportal.org/).

### External transcriptome data sets

Seven microarray data sets with transcriptome expression profiles and overall clinical outcomes were used for external validation. Patients with early-stage BCa were defined as samples with a T-stage of Ta or T1, indicating that these tumours were only in the innermost layer of the bladder lining (Ta) or had started to grow into the connective tissue beneath the bladder lining (T1); these tumours were commonly classified as NMIBC and were initially removed for this study. Specifically, five microarray data sets that were quantified by Illumina beadchip were combined as the MIBC-ILLUMINA cohort, including GSE13507 (n = 61) [10], GSE32548 (n = 38) [11], GSE32894 (n = 51) [2], GSE48075 (n = 72) [12], GSE48276 (n = 64) [3]; two microarray data sets that were sequenced by Affymetrix genechip were combined as the MIBC-AFFY cohort with the removal of patients who received chemo/radiotherapies, including GSE31684 (n = 74) [13] and E-MTAB-1803 (n = 43) [14]. Information about the platform and corresponding sample size of the eight MIBC data sets are summarized in Supplementary Table S1; demographic and clinical characteristic descriptions are detailed in Supplementary Table S2. For microarray data, the median value was considered if the gene symbol was annotated with multiple probe IDs. The potential cross-dataset batch effect was removed under an empirical Bayes framework by the R package “*sva*” [15], and the batch effect was further investigated using principal component analysis (Supplementary Figure S1a-b).

### Multi-omics integration and visualization

To perform integrative clustering, the MIBC-TCGA multi-omics data sets were primed to form five data matrixes of which rows correspond to the features and columns correspond to the common samples (n = 396). The transcriptome expression profile was first log_2_ transformed. For the methylation data, we extracted probes located in promoter CpG islands, and the median β value was considered for genes having more than one probe mapping to its promoter, resulting in 10,871 methylated genes. For the mutation matrix, a gene was considered mutated (entry of 1) if it contained at least one type of the following nonsynonymous variations: missense/nonsense/nonstop mutation, frameshift deletion/insertion, in-frame deletion/insertion, translation start site or splice site mutation; otherwise, 0 was used to designate wild-type status. For the CNAs, we condensed the genomic segments as described in the literature [16]. To better fit the model and accelerate the clustering efficiency, features with flat values were removed. Specifically, we selected the top 1,500 most variable mRNAs, lncRNAs, methylation genes, and copy number regions according to the median absolute deviation. Additionally, genes with mutation rates >3% (n = 1,358) were selected for subtyping. To find an optimal clustering number, we referred to the number of previous molecular subtypes of MIBC and calculated the clustering prediction index (CPI) and gap statistics [17]. Consequently, integrative clustering of the MIBC-TCGA cohort was independently conducted by 10 state-of-the-art multi-omics integrative clustering algorithms [8]. We borrowed the idea of a consensus ensemble for later integration of the clustering results derived from different algorithms to improve the clustering robustness [18].

### Calculation of microenvironment cell abundance and pathway enrichment

To establish a compendium of gene list related to specific microenvironment cells, two gene signatures (CIBERSORT [19] and MCPcounter [20]) were modified. As CIBERSORT does not contain signatures related to fibroblasts and endothelial cells, extra 40 genes were added to account for these cells (32 genes for endothelial cells and 8 genes for fibroblasts) from MCPcounter to our compendium, which consisted of 364 genes representing 24 microenvironment cell types. We then used gene set variation analysis (GSVA) on these gene sets to generate enrichment scores for each cell using the R package “*GSVA*”. The presence of infiltrating immune/stromal cells in the tumour tissue was estimated by the R package “*ESTIMATE*” [21]. Furthermore, the score of DNA methylation of tumour-infiltrating lymphocyte (MeTIL) in the MIBC-TCGA cohort was calculated individually according to the protocols outlined in the literature [22]. We referred to a published paper and the angiogenesis gene set(https://www.gsea-msigdb.org/gsea/msigdb/cards/ANGIOGENESIS) to construct a signature of ten oncogenic pathways, and the GSVA method was harnessed to generate enrichment scores [23]. A total of 21 replication stress signatures were retrieved from the literature, and single-sample gene set enrichment analysis (ssGSEA) was performed to quantify the enrichment level [24]. Two subtypes of replication stress were identified by hierarchical clustering.

### Bioinformatic analyses

We analysed the mutation landscape by the R package “*maftools*” with the initial removal of 100 FLAGS [25], and we evaluated the mutational signatures through the R package “*deconstructSigs*” [26]. Recurrent focal somatic CNAs were detected and localized by GISTIC2.0 through GenePattern(https://www.genepattern.org/), with the thresholds of copy number amplifications/deletions being equal to ± 0.3 (q-value < 0.05) [27]. The individual fraction of copy number-altered genome (FGA) for the MIBC-TCGA cohort was calculated based on copy number segment data as follows:

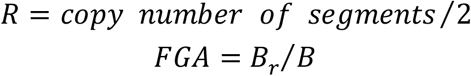

The FGA is the fraction of the genome with an value of log_2_(copy number) larger than 0.3 versus the genome with copy number profiled where *B*_*r*_ denotes the number of bases in segments with |*log*_2_*R*| > 0.3 and *B* represents the number of bases in all segments [17]. Differential methylation analysis was conducted on probes located in promoter CpG islands between tumour and adjacent normal samples by the R package “*ChAMP*” [28]; hypermethylated promoters were determined according to the following stringent criteria: the mean methylation β value in the tumour samples was greater than 0.5 and less than 0.2 in the adjacent normal samples with FDR < 0.05. Except for Lund [2], one nearest neighbour (oneNN) prediction model-based [3] and prediction analysis of microarray-based (PAM)[4] subtypes that were previously identified for MIBC-TCGA, we used the R package “*consensusMIBC*” to broadly predict individual consensus molecular subtypes (CMSs) for each MIBC, including basal/squamous (Ba/Sq), luminal papillary (LumP), luminal unstable (LumU), luminal non-specified (Luminal), neuroendocrine-like (NE-like), and stroma-rich subtypes [6]. Each sample in the external cohorts was further classified as one of the identified iCSs by nearest template prediction (NTP)[29].

### Regulon analysis

As previously described[5], we used the R package “*RTN*” to reconstruct transcriptional regulatory networks (regulons) including a total of 23 transcription factors (TFs) that were associated induced/repressed targets; another panel of 71 candidate regulators that were relevant to cancerous chromatin remodelling was also investigated [30]. Specifically, mutual information analysis and Spearman rank-order correlation deduced the possible associations between a regulator and all potential target from the transcriptome expression profile, and permutation analysis was utilized to erase associations with an FDR > 0.00001. Bootstrapping strategy removed unstable associations through one thousand times of resampling with consensus bootstrap greater than 95%. Data processing inequality filtering eliminated the weakest associations in triangles of two regulators and common targets. Individual regulon activity was estimated by two-sided GSEA.

### Therapeutic response analyses

Based on the drug sensitivity and phenotype data from GDSC 2016 (https://www.cancerrxgene.org/), the R package “*pRRophetic*” was employed to predict the chemotherapeutic sensitivity for each MIBC sample using the expression profiles of 727 human cancer cell lines (CCLs) as the training cohort; the IC_50_ (lower IC_50_ indicates increased sensitivity to treatment) of each sample treated with a specific chemotherapeutic agent was estimated by ridge regression, and 10-fold cross-validation was used to measure the prediction accuracy [31]. For immunotherapy, we harnessed subclass mapping to infer the clinical response to immune checkpoint inhibitors [32]. In this manner, we first retrieved a published data set consisting of 47 melanoma patients who responded to immunotherapies [33]. We then downloaded the IMvigor210 transcriptome profile (n = 298), which originated from a phase II trial that investigated the clinical activity of atezolizumab (anti-PD-L1 agent) for locally advanced and metastatic urothelial carcinoma (mMIBC)[34]. We calculated the transcriptome TPM values and extracted the best confirmed overall response information using the R package “*IMvigor210CoreBiologies*” [35]. We collated a list of immune-related genes (IRGs) from the nCounter PanCancer Immune Profiling Panel that consisted of 770 unique genes closely associated with the human immune response [36]; transcriptome expression of 754 matched IRGs was extracted for subclass analysis.

### Statistical analyses

All statistical tests were conducted by R4.0.2, including two-sample Mann-Whitney test for continuous data, Fisher’s exact test for categorical data, log-rank test for Kaplan-Meier curves, and Cox proportional hazards regression for estimating the hazard ratios (HRs) and 95% confidence interval (CI). The treatment effect of immune checkpoint inhibitors was measured by two nonproportional hazards statistical approaches, namely, restricted mean survival (RMS) and long-term survival inference, by using the R packages “*survRM2*” and “*ComparisonSurv*”, respectively [37]. A random forest (RF) predictive model was developed by the R package “*varSelRF*”, technical details of which have been described previously [38]. The receiver operating characteristic (ROC) curves were used to assess the model predictive performance. Most of the above analytic processes are embedded in the R package “*MOVICS*”, which we recently developed for multi-omics integration and visualization [17]. For all unadjusted comparisons, a two-tailed *P* < 0.05 was considered statistically significant.

## RESULTS

### Multi-omics integrative molecular subtypes of MIBC

We determined the optimal cluster number of four taking into account two clustering statistics and previous molecular classifications (Supplementary Figure S2a). A consensus ensemble generated from ten multi-omics integrative clustering approaches identified four robust iCSs (Supplementary Figure S2b), which presented with distinctive molecular patterns across transcriptome expression, epigenetic methylation, CNA and somatic mutation (Figure 1a). These classifications were significantly associated with age, sex, pathological stage and previously identified molecular classifications (Supplementary Table S3). As most molecular subtypes of MIBC were defined through gene expression which may directly alter biological functions, the iCSs were compared with those gene expression-based classifications; we found that iCS1 and iCS4 significantly overlapped with the basal-like subtypes (*P* < 0.001), whereas iCS2 and iCS3 were enriched for the luminal-like subtypes (*P* < 0.001; Figure 1b). We then relabelled iCS1 through iCS4 as iBS1, iLS2, iLS3, and iBS4, respectively. Interestingly, iBS1 and iBS4 were dramatically distinguished from the previously defined basal-like subtype, which caught our attention. Our classification system was tightly associated with PFS (*P* = 0.056; Figure 1c) and OS (*P* < 0.001; Figure 1d) rates. Strikingly, iBS4 showed the worst overall survival rate among all four iCSs (all pairwise comparisons, *P* < 0.05; Figure 1d).

**Figure 1.**
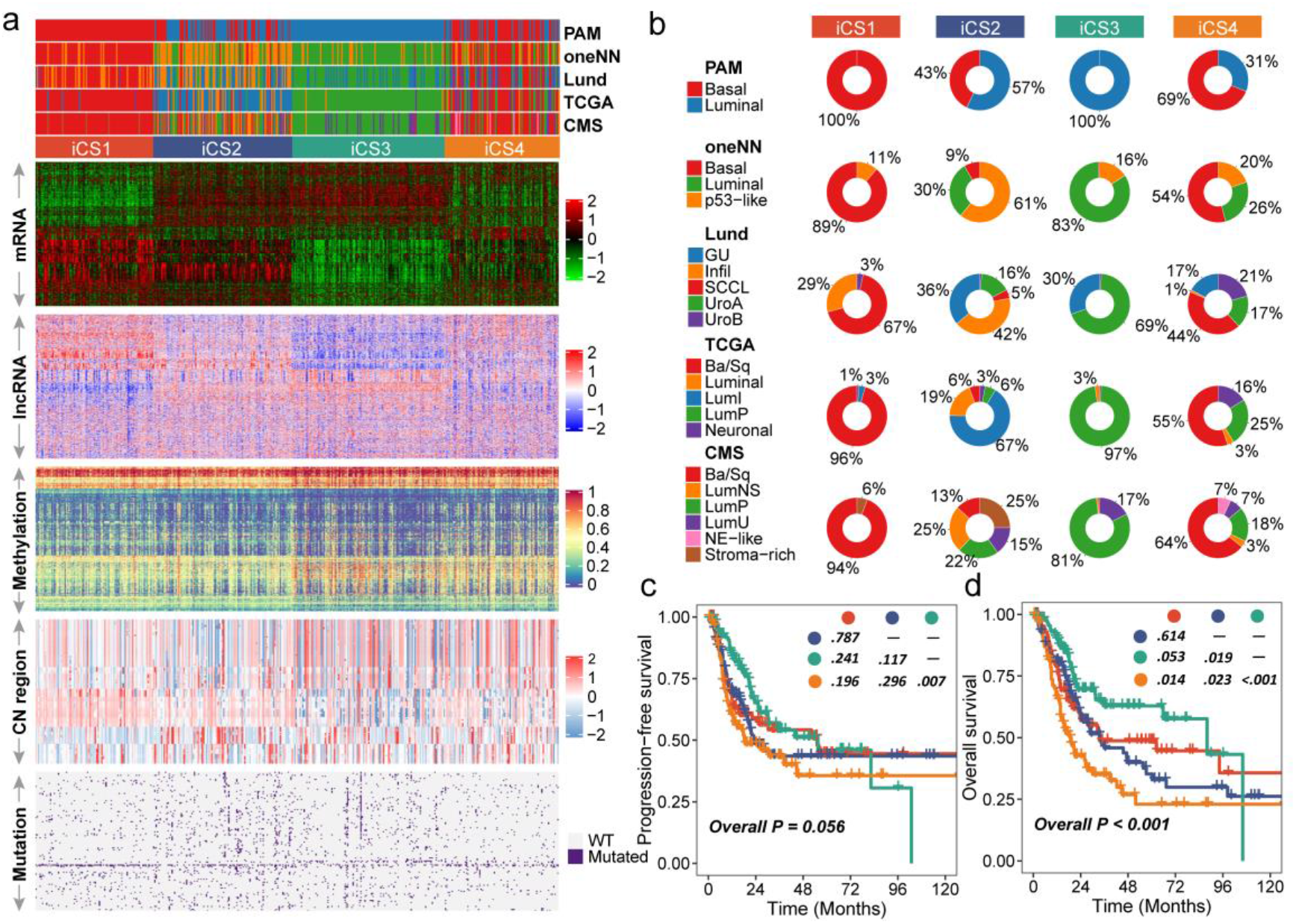
The multi-omics consensus ensemble identifies four molecular subtypes of MIBC. a) Comprehensive heatmap showing the molecular landscape of four integrative consensus subtypes (iCSs) of MIBC (n = 396). Other previously defined gene expression-based MIBC subtypes were annotated at the top of the heatmap, including prediction analysis of microarrays-based (PAM), one nearest neighbour (oneNN) prediction model-based, Lund, TCGA and consensus molecular subtypes (CMS). b) Pie charts showing the proportion of other gene expression-based MIBC subtypes in the current iCS. Kaplan-Meier curves of overall survival and progression-free survival with log-rank test for 396 MIBC patients stratified by iCS.

### Delineation of integrative consensus subtypes of MIBC

To further explore transcriptomic differences, we analysed regulons for 23 MIBC-specific TFs and potential regulators relevant to cancerous chromatin remodelling [5,30], leading to a strong confirmation of the biological pertinency of the four-classification because the regulon activity was tightly associated with iCSs (Figure 2a). Similar patterns of regulon activity were shared by iBS1 and iBS4, but iBS1 differed with high activity of *ESR1, FGFR1*, and *GATA6*, while iBS4 was distinctly associated with high activity of *TP63*. Regulons of *AR, GATA3, PPARG, ERBB2*, and *ESR2* were significantly activated in iLS2 and iLS3, whereas *FGFR3, RARG, RXRA, RXRB, ERBB3* and *FOXA1A* were uniquely enriched in iLS3. Regulon activity profiles that were associated with cancerous chromatin remodelling highlighted other possible differential regulatory patterns among four iCSs, indicating that epigenetically driven transcriptional networks might be important differentiators of these molecular subtypes (Figure 2a). The potential epigenetic differences among the subtypes might be further supported by differential methylation analysis, which demonstrated that iLS3 (265 vs. 45 in iLS2) and iBS4 (191 vs. 26 in iBS1) had more hypermethylated promoters (296 unique loci) than the 21 adjacent normal bladder samples had (Figure 2a).

**Figure 2.**
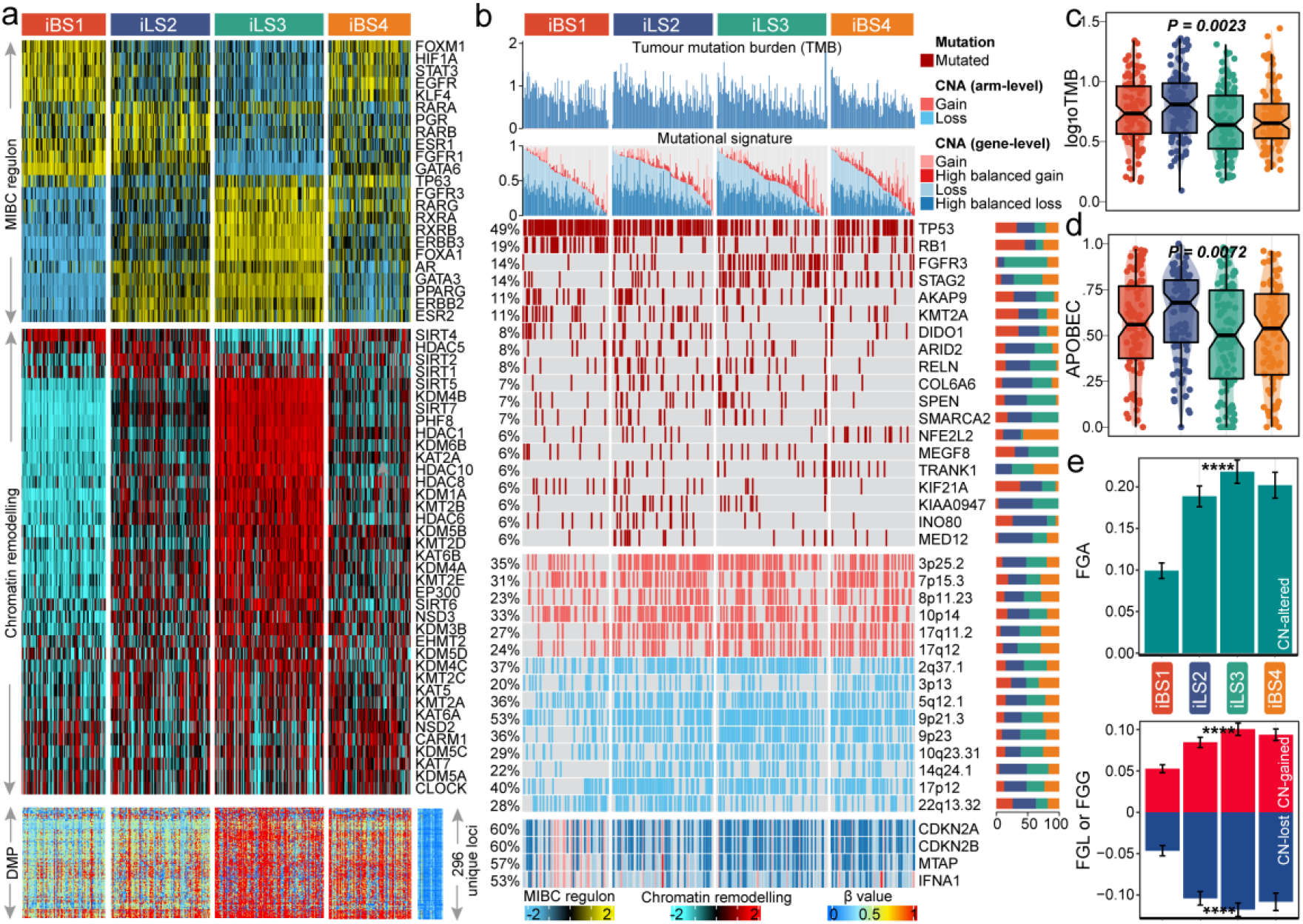
Molecular landscape of four MIBC iCSs. a) Heatmap showing regulon activity profiles for 23 transcription factors (top panel), potential regulators associated with chromatin remodelling (middle panel), and 296 unique differentially methylated promoters derived from each iCS vs. adjacent normal samples (bottom panel). b) Genomic alteration landscape according to iCS. Samples are ordered by the combined contribution of APOBEC-related mutational signatures (SBS2 + SBS13) with each iCS. Tumour mutation burden (TMB), relative contribution of four mutational signatures, selected differentially mutated genes (>5%) and broad-level copy number alterations (>20%), and selected genes located within chromosome 9p21.3 are shown from the top to the bottom panels. The proportion of iCS in each alteration is presented in the right bar charts. The distributions of TMB and APOBEC contributions are shown in c) and d), respectively. e) Distribution of fraction genome altered (FGA) and fraction genome gain/loss (FGA/FGG). Bar charts are presented as the mean ± standard error of the mean.

To investigate the genomic heterogeneity of MIBC further, we found that iLS2 showed a significantly higher tumour mutation burden (TMB, *P* = 0.002; Figure 2b-c) than the other subtypes. We inferred that four mutational signatures showed a high correlation with bladder cancer, namely, SBS1 (age-related), SBS2 and SBS13 (APOBEC activity-related) and SBS5 (*ERCC2* mutation-related). We found that iLS2 had significantly more mutations in APOBEC-related signatures (*P* = 0.007; Figure 2b and d). Of the frequently mutated genes (>5%), iBS1 harboured significantly more mutations of *TP53* (74%; *P* < 0.001), *RB1* (39%; *P* < 0.001) and *KMT2A* (18%; *P* = 0.033) than the other subtypes, while iBS4 was significantly enriched in mutations of *NFE2L2* (also known as *NRF2*) (16%; *P* = 0.001; *P* = 0.005 compared to iBS1 [3%]) and *TRANK1* (10%; *P* = 0.06; *P* = 0.001 compared to iBS1 [0%]); *KIAA0947* (11%; *P* = 0.005), *MED12* (11%; *P* = 0.005), *COL6A6* (13%; *P* = 0.008), and *ARID2* (14%; *P* = 0.01) were mutated more frequently in iLS2, whereas iLS3 was significantly enriched in mutations of *FGFR3* (34%; *P* < 0.001), *STAG2* (22%; *P* = 0.006), and *SPEN* (11%; *P* = 0.05) (Figure 2b; Supplementary Table S4).

We then investigated chromosomal instability by calculating the FGA scores and found that iBS1 had better chromosomal stability than the other subtypes with significantly lower copy number loss or gain (both, *P* < 0.001; Figure 2e). Human chromosome 9p21.3 is susceptible to inactivation in cell immortalization and diseases, such as cancer [39]; within this region, we found that interferon alpha (*IFNA*) genes, *MTAP* and *CDKN2A/B* were differentially altered in iBS1 compared to the other subtypes, which may contribute to shaping different basal-like subtypes (Figure 2b).

### Prognostic value of subtype-specific signatures in external MIBC cohorts

Given that transcriptome-level data were the most commonly used molecular profiles in cancer research, we identified 30 mRNAs with uniquely and significantly upregulated expression as classifiers for each subtype in the MIBC-TCGA cohort, and a 120-gene signature was generated to predict the identified MIBC subtypes in the external data sets individually (Figure 3a; Supplementary Table S5). NTP classified each sample in the external cohorts as one of the identified iCSs (Figure 3b-c). The signature-predicted iCSs highly overlapped with CMS but further refined the basal-like subtype classification (both, *P* < 0.001; Figure 3d-e). Consistently, iBS4 in the MIBC-ILLUMINA cohort had the most unfavourable prognosis out of all the subtypes (all pairwise comparisons, *P* < 0.005; Figure 3f); similar results were found in the MIBC-AFFY cohort (Figure 3g).

**Figure 3.**
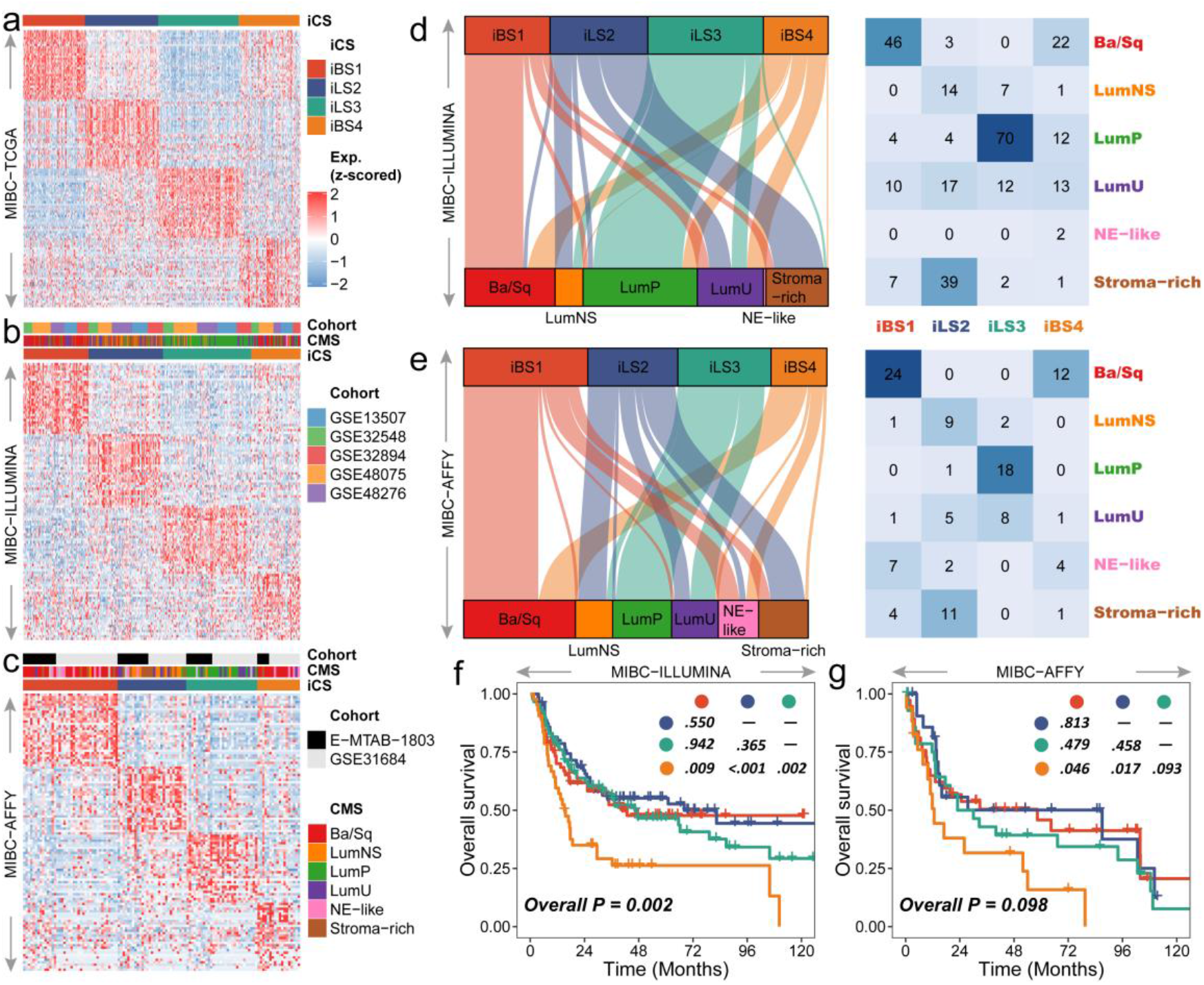
Validation of the 120-gene signature to reproduce four MIBC iCS in external cohorts. Heatmap showing the transcriptome expression pattern of the 120-gene signature in nearest template predicted iCS of three MIBC cohorts, including a) MIBC-TCGA, b) MIBC-ILLUMINA and c) MIBC-AFFY. Overlap between iCS and consensus molecular subtype (CMS) is represented in d) for MIBC-ILLUMINA and e) for MIBC-AFFY. Several patients in the MIBC-AFFY cohort had no predicted CMS because the samples showed Pearson’s correlation between the gene expression profile and consensus centroid profiles lower than the confidence minimal threshold of 0.1. Kaplan-Meier curves of overall survival with the log-rank test for MIBC patients stratified by iCS are shown in f) for MIBC-ILLUMINA and g) for MIBC-AFFY.

### Differential immune profiles across MIBC subtypes

Since cancer immunity plays a critical role in tumour progression, we suspected that the tumour microenvironment of the iBS4 subtype may be different from that of the iBS1 subtype. Thus, we investigated the specific immune cell infiltration status of samples from three cohorts; we quantified the infiltration levels of 24 microenvironment cell types and surveyed the MIBC samples for the expression of immune checkpoints. The analysis of gene expression signatures suggested that immunocyte infiltration was dramatically higher in both the iBS1 and the iLS2 subtype, while it was relatively lower in abundance in the iBS4 subtype and was lowest in the iLS3 subtype (Figure 4a). Compared to the other subtypes, iBS1 had relatively higher expression of several genes that represent potential targets for immunotherapy (Figure 4a), including *PDCD1* (PD1), *CD247* (CD3), *CD274* (PDL1), *PDCD1LG2* (PDL2), *CTLA4* (CD152), *TNFRSF9* (CD137), *TNFRSF4* (CD134) and *TLR9*. The landscape of immune profiles was validated in both the MIBC-ILLUMINA and MIBC-AFFY cohorts (Supplementary Figure S3a-b). Additionally, the iBS1 subtype in the MIBC-TCGA cohort had a significantly higher MeTIL score than other iCSs, indicating a higher fraction of tumour-infiltrating leukocytes (*P* < 0.001, Figure 4a). Given that the iBS1 subtype has a relatively higher mutational load and immune cell infiltration level as well as a lower chromosomal instability than the other subtype, this subtype might also have a high level of neoantigens.

**Figure 4.**
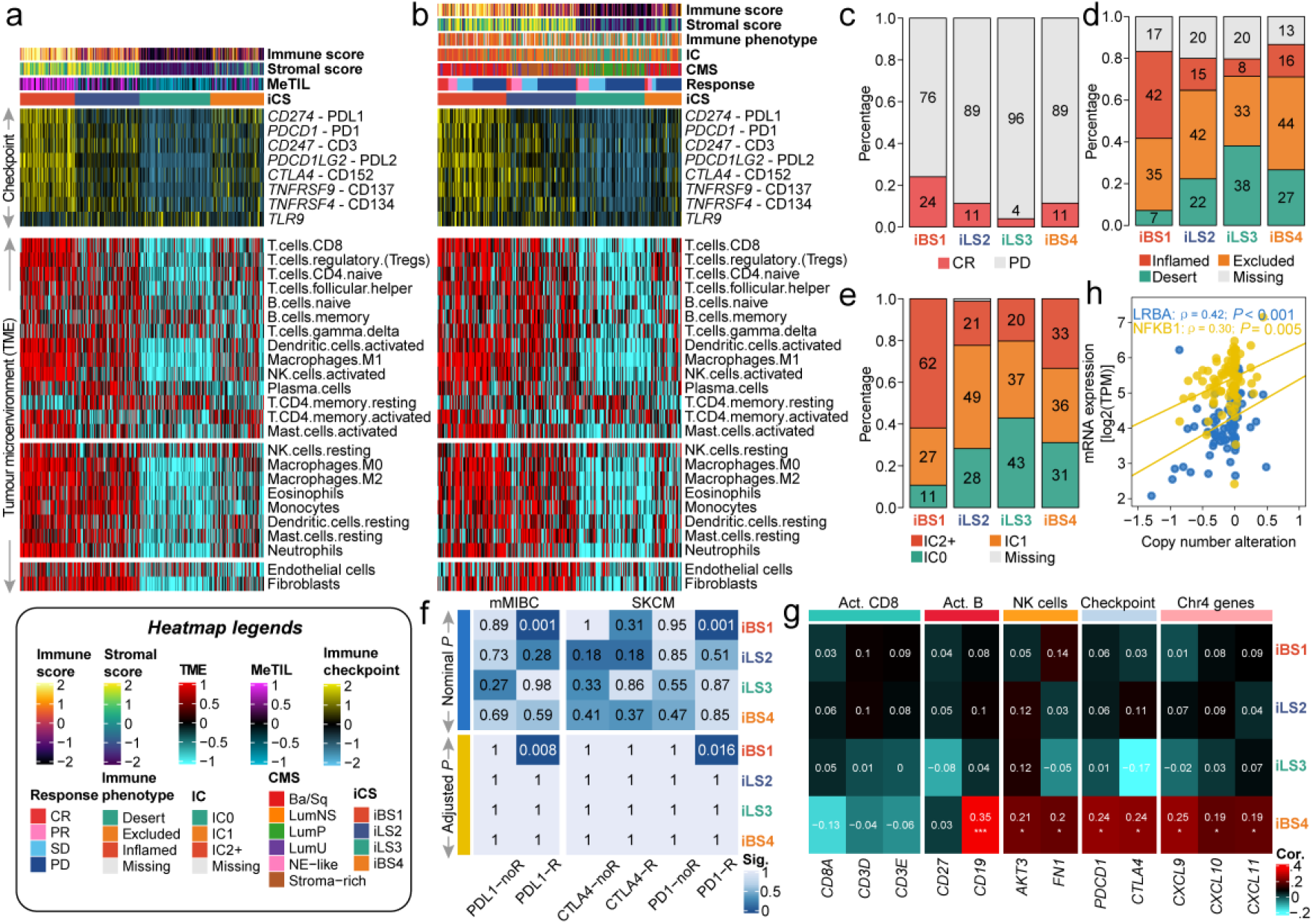
Differential immune profile across MIBC subtypes and its association with genomic alteration and immunotherapeutic response. a) Heatmap showing the immune profile in the MIBC-TCGA cohort, with the top panel showing the expression of genes involved in immune checkpoint targets and the bottom panel showing the enrichment level of 24 microenvironment cell types. The immune enrichment score, stromal enrichment score and DNA methylation of tumour-infiltrating lymphocytes (MeTILs) were annotated at the top of the heatmap. b) Immune profile heatmap for the IMvigor210 cohort with annotations for immune enrichment score, stromal enrichment score, immune phenotype, immune cell (IC) level, and best confirmed overall response, including progressive disease (PD), stable disease (SD), partial response (PR) and complete response (CR). Barplots showing the distribution of c) CR and PD, d) immune phenotype and e) IC levels in iCS of the IMvigor210 cohort. f) Subclass analysis revealed that the iBS1 subtype could be more sensitive to the anti-PD-L1 and anti-PD1 agents (both, Bonferroni-corrected *P* = 0.001) using two reference cohorts in which patients received immunotherapy. g) Heatmap showing the correlation between chromosome 4 copy number deletion and expression profiles of immune markers across ICSs in the MIBC-TCGA cohort. A positive correlation indicates that the deletion of chromosome 4 may trigger the downregulation of the relevant immune gene expression. h) Positive correlation between downregulation of expression of immune regulatory genes and chromosome 4 deletion in immune-cold iBS4 of the TCGA-MIBC cohort.

To investigate whether the identified immune profiles could be reproduced in MIBC patients who received pretreatment, we analyzed RNA-seq data from the IMvigor210 cohort (n = 298) in which the clinical effect of PD-L1 blockade with atezolizumab was evaluated in mMIBC patients [34]; we found a similar immunologic landscape (Figure 4b). Additionally, a significantly higher proportion of patients with iBS1-subtype tumours achieved a complete response with atezolizumab treatment than patients in with iCS tumours (*P* = 0.024; Figure 4c). To evaluate the treatment effect of immune checkpoint inhibitors on different iCSs in the IMvigor210 cohort, we compared the RMS time difference at six months and one year after treatment. We found that the iBS4 subtype showed a significantly poorer outcome than immune-hot iCSs (*i*.*e*., iBS1 and iLS2; both *P* < 0.05 at 6 months, both *P* < 0.1 at 12 months) and had a segregated survival curve compared to the immune-cold iLS3 subtype (Supplementary Figure S4a). Due to the delayed clinical effect of immunotherapy, we also compared the long-term survival rates after three months of treatment. Consistently, the iBS4 subtype was associated with significantly poorer long-term survival than the immune-hot iBS1 (*P* = 0.046) and iLS2 (*P* = 0.027) subtypes and the immune-cold iLS3 subtype (*P* = 0.095), suggesting its high malignancy and potential resistance to immune checkpoint blockade (Supplementary Figure S4b). Mariathasan *et al*. reported that the clinical effect of anti-PD-L1 blockade may be influenced by the tumour-immune phenotype and immune cell (IC) subtypes [40]. We found that 42% of patients with iBS1-subtype tumours were defined as having an “inflamed” immunophenotype (*P* < 0.001) and enriched IC2+ subtype (62%, *P* < 0.001). Compared to the iBS1 subtype, the iBS4 subtype was significantly enriched in the “noninflamed” immunophenotype (*P* = 0.001) category and showed a relatively equal distribution in the IC subtype classification. Forty-two percent of patients with iLS2-subtype tumours were defined as having an “excluded” immunophenotype and an enriched IC1 subtype (*P* = 0.003), whereas the iLS3 subtype was enriched in the the “desert” immunophenotype (*P* < 0.001) and the IC0 subtype (*P* < 0.001) (Figure 4d-e). Thus, we renamed the iBS1 subtype “basal-inflamed”, the iBS4 subtype “basal-noninflamed”, the iLS2 subtype “luminal-excluded”, and the iLS3 subtype “luminal-desert”.

### The basal-inflamed subtype has a high likelihood of responding to immunotherapy

Although three PD-L1 blockades (atezolizumab, durvalumab, and avelumab) and two PD-1 blockades (pembrolizumab and nivolumab) have been approved by the Food and Drug Administration for the adjuvant treatment of BCa, only a subgroup of patients respond to these immunotherapies. Considering the dramatically diverse tumour microenvironments, we next investigated whether there is a difference among MIBC subtypes in the likelihood of responding to immune checkpoint blockade. To this end, we performed subclass mapping of all three MIBC cohorts and revealed that only the basal-inflamed MIBC subtype showed highly similar immune profiles to the patients in the IMvigor210 cohort who responded to PD-L1 checkpoint inhibitors (all, FDR < 0.01; Figure 4f, Supplementary Figure S5). Additionally, the basal-inflamed MIBC subtype had transcriptome-level similarity to a subgroup of melanoma patients who responded to anti-PD1 blockade (all, FDR < 0.05; Figure 4f, Supplementary Figure S5), which indicated that the current classification may be useful to identify ideal candidates for immunotherapy, especially for basal-like MIBC.

### Association between chromosome 4 deletion and the MIBC tumour microenvironment

Recently, Hao *et al*. suggested that chromosome 4 (Chr4) loss could induce an unfavourable “cold” tumour immune microenvironment [41]. We then decided to investigate whether such an association between CNA and immunophenotype existed in the MIBC-TCGA cohort. We performed GISTIC2.0 for each iCS and found that only the basal-noninflamed subtype showed significant Chr4p and 4q deletions (FDR = 0.06 for 4p and 0.17 for 4q; Supplementary Table S6). Chr4 harbours several immune regulatory genes (*e*.*g*., *FKB1* and *LRBA*), and genes that encodes chemokines (*e*.*g*., *CXCL9*/*10*/*11*) which are crucial for T cell recruitment. Additionally, deletion of Chr4 genetically linked with immune deficiency syndromes [42]. Integrative analysis of RNA-seq expression and CAN indicated that the decreased expression of key immune markers was tightly associated with Chr4 deletion in the basal-noninflamed subtype; these markers included immune checkpoint targets (*e*.*g*., *PDCD1* and *CTLA4*) and genes related to activated B cells, natural killer cells, and cytokines (*e*.*g*., *CXCL9*/*10*/*11*) that regulate immune cell activation and trafficking (Figure 4g), and their key mediator, *NFKB1*, was also significantly positively correlated with its copy number deletion in the basal-noninflamed subtype (Figure 4h). We then analyzed the association between Chr4 loss and prognosis in basal-like MIBC. Univariate analysis revealed that Chr4p loss was tightly associated with unfavourable prognosis (HR = 1.56, 95% CI: 1.001-2.437, *P* = 0.0496); we did not observe statistical significance regarding Chr4q loss (HR = 1.307, *P* > 0.05).

### Dysfunction of oncogenic pathways in MIBC

We then investigated the activation of oncogenic pathways among iCSs (Figure 5a; Supplementary Figure S6a). We found that the cell cycle oncogenic pathway was significantly activated in basal-inflamed/noninflamed MIBC (all, *P* < 0.001; Figure 5b, Supplementary Figure S6b). Activation of the cell cycle pathway activates cell cycle checkpoint regulatory proteins (*e*.*g*., ATR and WEE1) involved in replication stress (RS), which has been described to be closely associated with DNA damage responses that contribute to cisplatin resistance. Consequently, we investigated the potential of targeting RS as a therapeutic regimen for a subgroup of MIBCs. Unsupervised hierarchical clustering was performed by using 21 replication stress signatures in all three MIBC cohorts, resulting in two RS subtypes (*i*.*e*., RS-High and RS-Low) (Figure 5c; Supplementary Figure S6c). In each cohort, the basal-inflamed/noninflamed MIBC subtypes were significantly enriched in the RS-High subtype (all, *P* < 0.001), suggesting potential cisplatin-based chemoresistance. To test the potential therapeutic efficiency of cell cycle checkpoint inhibitors for the basal-inflamed/noninflamed MIBC subtypes, we estimated the IC_50_ for three ATR inhibitors (*i*.*e*., VE-821, VE-822 and AZD6738) and two WEE1 inhibitors (*i*.*e*., Wee1 inhibitor and MK-1775) available from GDSC (Version: 2016) through a predicted model-based strategy. Specifically, we constructed a ridge regression model based on 727 CCLs with available IC_50_ measurements for these five agents. In general, both RS-High and basal-inflamed/noninflamed MIBC subtypes in all three MIBC cohorts were more susceptible to cell cycle checkpoint inhibitors than other MIBC subtypes (Figure 5d-e; Supplementary Figure S6d).

**Figure 5.**
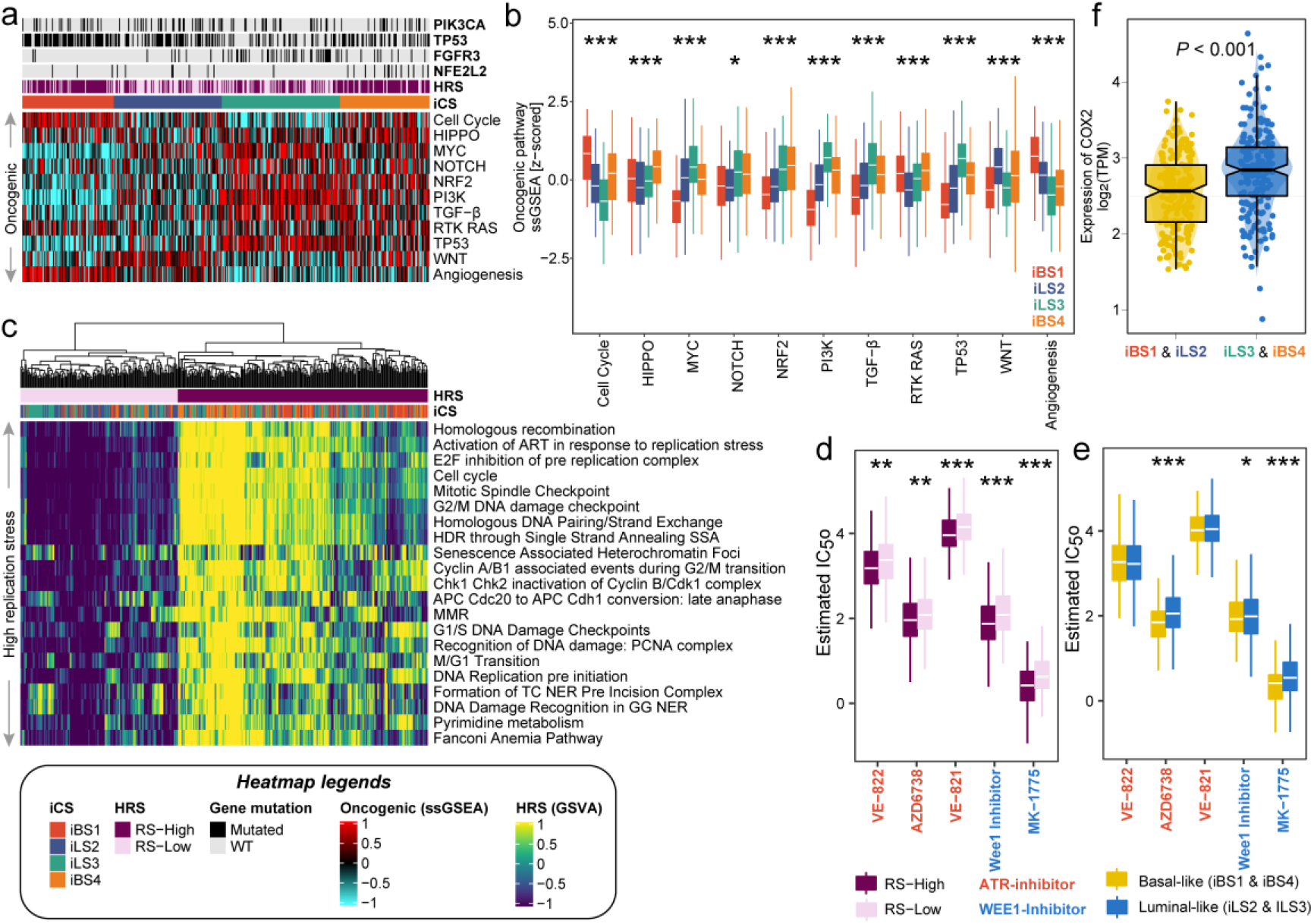
Oncogenic pathways, replication stress and targeted inhibitors in MIBC. Dysfunctional oncogenic pathways quantified by single-sample gene set enrichment analysis are presented in a) a heatmap and b) a boxplot for the TCGA-MIBC cohort. Relevant mutations involved in several oncogenic pathways are annotated at the top. c) Heatmap of pathways and molecular processes (Reactome database) involved in DNA maintenance and cell cycle regulation activated in replication stress and DNA damage response. Two replication stress (RS) subtypes were identified for the TCGA-MIBC cohort. Subtypes with d) high replication stress (RS-High) or e) basal-like MIBC (iBS and iBS4) were inferred to be much more sensitive to both ATR (*i*.*e*., VE-822, AZD6739 and VE-821) and WEE1 (*i*.*e*., Wee1 inhibitor and MK-1775) inhibitors by applying a ridge regression model using 727 human cancer cell lines. Drug sensitivity was measured as ln(IC_50_), and the lower the value was, the more sensitive the patient would be to the treatment. f) Distribution of *COX2* expression between immune-hot (*i*.*e*., iBS1 and iLS2) and immune-cold (*i*.*e*., iLS2 and iBS4) phenotypes of MIBC-TCGA.

Additionally, the luminal-desert and basal-noninflamed subtypes showed relatively higher activation of the oncogenic *NRF2* pathway; the luminal-excluded subtype was highly enriched in the *WNT* pathway, while the luminal-desert subtype showed the lowest enrichment in angiogenesis genes but the highest activation of the *TGF-β* pathway (Figure 5b). A recent study reported that *NRF2* enables an immune-cold environment by inducing *COX2*/*PGE2* and inhibiting the DNA-sensing innate immune response [43]. Thus, we assessed the expression level of *COX2* in the luminal-desert/basal-noninflamed subtypes and the other MIBC subtypes in cohorts where *COX2* expression data is available (*i*.*e*., the MIBC-TCGA and IMvigor210 cohorts). Consistently, the luminal-desert/basal-noninflamed subtypes had significantly higher *COX2* expression levels than the other MIBC subtypes in both the TCGA (*P* < 0.001; Figure 5f) and IMvigor210 (*P* = 0.083; Supplementary Figure S7a-c) cohorts, suggesting that dysfunction of oncogenic pathways might drive the low immune infiltration of these MIBCs.

### Random forest classifier to refine basal-like subtypes

Given the distinct molecular and prognostic characteristics of the basal-inflamed/noninflamed MIBC subtypes compared with the traditional basal-like MIBC classification, we developed a predictive model to refine the classification of basal-like subtypes using the RF algorithm. To this end, we performed differential expression analysis on ∼700 IRGs in basal-inflamed and basal-noninflamed MIBC in the TCGA cohort to select informative IRGs for the RF model (the top 50 IRGs ordered by log_2_FoldChange for each subtype with FDR < 0.05). The final basal classifier contains five IRGs, *C3AR1, CCL8, FCGR3A, LILRB2*, and *PDCD1LG2*; the predictive model achieved a prediction accuracy of 100% in the TCGA-MIBC cohort and 90.8% in the ILLUMINA-MIBC cohort, 85.3% in the AFFY-MIBC cohort and 92.0% in the IMvigor210 cohort (Figure 6a). Additionally, we demonstrated the capability of prognosis stratification when deploying the basal-classifier to refine the classification of basal-like subtypes, including TCGA-Ba/Sq (n= 139; *P* = 0.013), Lund-SCCL (n = 103; *P* = 0.066), PAM-basal (n = 194; *P* = 0.001), oneNN-basal (n = 136; *P* = 0.001) and CMS-Ba/Sq (n = 155; *P* < 0.001), into basal-inflamed and basal-noninflamed subtypes (Figure 6b-f). We then investigated whether our basal classifier is capable of refining the classification of basal-like tumours with distinct immune profiles. Therefore, we conducted a pan-carcinoma classification based on the previously identified basal-like subtype across 22 different carcinomas (n = 2,459) [44]. Strikingly, almost all kinds of epithelial tumours can be refined into immune-hot or immune-cold basal-like phenotypes (Figure 6g). The RF basal-classifier and the NTP 120-gene signature were merged into the R package “*refineMIBC*”, which is documented and freely available at https://github.com/xlucpu/refineMIBC.

**Figure 6.**
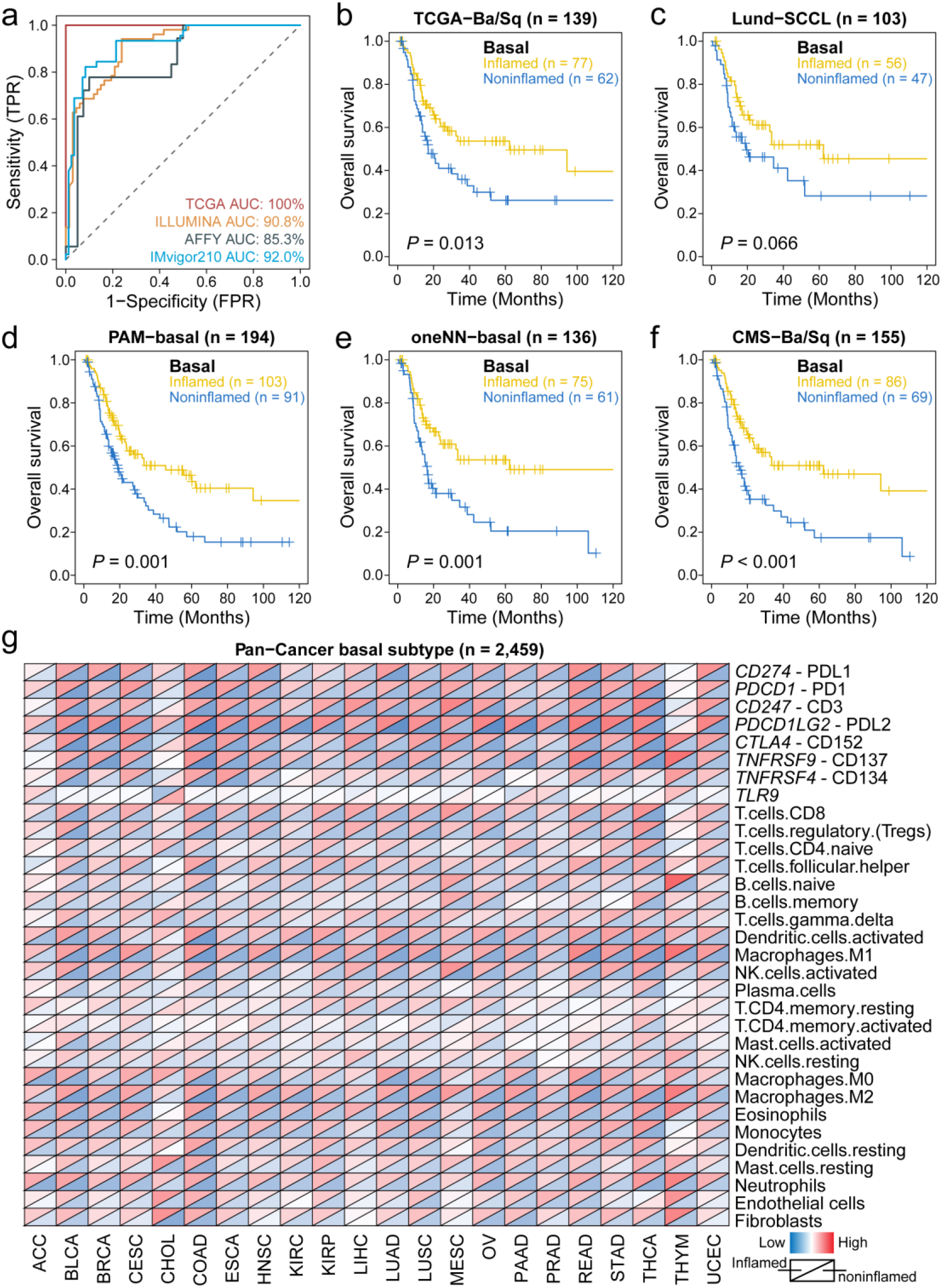
Predictive performance of random forest basal-classifier and its application in pan-carcinoma investigation. a) ROC curve showing prediction accuracy when using the basal-classifier to refine basal-like MIBC into basal-inflamed and basal-noninflamed subtypes. The prognostic value of the basal-classifier in refining previously identified basal-like subtypes of MIBC is presented in b) for the TCGA-basal/squamous subtype, c) for the Lund-SCCL subtype, d) for the PAM-basal subtype, e) for the oneNN-basal subtype, and f) for the CMS-Ba/Sq subtype. g) Diagonal heatmap showing global immunological divergence across 22 basal-like carcinomas with a total of 2,459 cases. The upper triangle of each heatmap cell represents the average expression of the immune checkpoint/cell in the predicted basal-inflamed subtype given a specific tumour type, while the lower triangle represents the predicted basal-noninflamed subtype.

## DISCUSSION

To refine classifications of MIBC and clinically relevant biomarkers, we conducted an integrative consensus ensemble analysis of MIBC under multi-omics framework and defined four iCSs, which shed new light on patient stratification for frontline therapies. We developed and validated a single-sample 120-gene signature to refine the classification of MIBC, and notably, we further built a 5-gene basal-classifier that is capable of distinguishing patients with an immune-cold phenotype and poor outcome from those with traditional basal-like MIBC. The 120-gene signature was successfully validated in three external cohorts with a total of 701 patients with data analyzed on different sequencing platforms, and the 5-gene classifier demonstrated robust predictive performance in addition to basal-like MIBC stratification, even for other basal-like epithelial tumours, suggesting a global immunological divergence across basal-like carcinomas, which might help identify ideal candidates for immunotherapy. Although the current subtypes were characterized by distinct molecular patterns and multi-omics perspectives should be incorporated into future classification paradigm for MIBC with larger cohorts, we suggest the utilization of the transcriptome-based signature/classifier for clinical application, as the expression profile could directly reflect tumour biological processes and possible treatment options.

Specifically, alterations involving *RB1* and *NFE2L2* are known to be enriched in basal-like MIBC [7, 45], but our study further revealed that the basal-inflamed MIBC subtype had more *RB1* mutations, whereas the basal-noninflamed MIBC subtype harboured a significantly higher number of *NFE2L2* (*NRF2*) mutations. Accordingly, we observed activation of the *NRF2* oncogenic pathway in the basal-noninflamed MIBC subtype, as well as elevated expression of *COX2*. A previous study showed that *NRF2* activation elevates the production of *COX2* and *PGE2*, thus suppressing the innate immune response and contributing to forming an immune-cold tumour environment [43]. Unfortunately, it is not feasible to combine *NRF2* inhibitors, which are currently unavailable, with checkpoint inhibitors. Nevertheless, preclinical models indicated that *COX* inhibitors synergize with anti-PD-1 immunotherapy [46]; thus, targeting the downstream markers of *NRF2* poses a therapeutic opportunity for immune-cold MIBCs. Additionally, we identified that the loss of Chr4 is associated with the immune-cold phenotype of the basal-noninflamed MIBC; combined with the activated *NRF2* pathway, this finding may shape understanding of an immune-cold subgroup of basal-like MIBC with unfavourable prognosis.

Recent studies showed tight connections between tumour genomic landscape and anti-tumour immunity. Particularly, the presence of neoantigens and total mutational burden may trigger responses of T cells, while tumour aneuploidy is associated with immune evasion and weakened sensitivity to immunotherapy [47]. In the current classification scheme, the basal-inflamed MIBC subtype had a relatively high mutation load but a widely stable genome. The highly infiltrated immune microenvironment and highly expressed checkpoint target genes enhance the likelihood of this subtype responding to immune checkpoint inhibitors. Nevertheless, the relatively immune-hot luminal-excluded MIBC subtype had dramatically high genomic instability and low expression of checkpoint genes, suggesting that these tumours would not be susceptible to immunotherapy. As expected, the basal-inflamed MIBC subtype was more sensitive to PD-L1 inhibitors than the other MIBC subtypes in the IMvigor210 cohort, whereas the basal-noninflamed MIBC presented with the shortest OS after treatment, indicating its high resistance to immune checkpoint blockade. The luminal-desert MIBC subtype was significantly enriched in *TGF-β* signalling genes, and activated *NRF2* pathways may synergistically form an immune-cold phenotype and attenuate the response to immunotherapy [40].

The different biological characteristics of the four subtypes may indicate the need for different therapeutic interventions. Of note, the activated cell cycle pathway in the basal-like MIBC subtype may lead to potential resistance to cisplatin, but this subtype may be susceptible to ATR and WEE1 inhibitors. The highly activated angiogenesis pathway in the basal-inflamed and luminal-excluded MIBC subtypes may increase sensitivity to *EGFR* inhibitors, and a phase II trial demonstrated anti-tumour activity of anti-*EGFR* agents [48]. The luminal-excluded and luminal-desert MIBC subtypes were characterized by activated *WNT* signalling and *FGFR3* mutations, respectively, suggesting that *WNT* and *FGFR* inhibitors could be effective for these tumours. Inhibitors of *FGFR* represent an outstanding treatment approach for luminal-papillary MIBC [49], which highly overlaps with luminal-desert MIBC. Given that a number of *Wnt/β-catenin* signalling inhibitors are being used for various human cancers [50], it is promising that such inhibitors could be developed as potential therapeutic candidates for *WNT* signalling-activated MIBC (*e*.*g*., luminal-excluded MIBC).

Several limitations should be acknowledged. First, the included cohorts varied in size, composition, and sequencing platform. Second, incomplete treatment information may have caused bias in the study design. Third, bulk RNA-seq and microarray profiles are confounded by signals quantified from a mixture of cell populations; thus, it is warranted in the future to combine these RNA-based discoveries with multiplex immunohistochemistry to delve intrinsic tumour cell alterations and their interplay with the tumour microenvironment that dictate therapy response.

Briefly, we identified four MIBC subtypes with distinct landscapes using a multi-omics approach that stratify prognosis, tumour microenvironment characteristics and distinct sensitivity to frontline therapies. We offer the R package “*refineMIBC*” as a research tool to refine the classification of MIBC from a single-sample perspective in retrospective or prospective studies. We expect this multi-omics consensus ensemble approach to MIBC classification will further assist precision medicine by providing a blueprint for the clinical development of rational targeted and immunotherapeutic strategies.

## DECLARATIONS

### Ethics approval and consent to participate

As the data used in this study are publicly available, no ethical approval was required.

## Consent for publication

Not applicable.

## Availability of data and material

The raw data for this study were generated at the corresponding archives. Derived data supporting the findings are available from the corresponding author [F.Y.] on reasonable request. We developed the R package, “*refineMIBC*”, which is documented and freely available at https://github.com/xlucpu/refineMIBC. This package implements a 120-gene template that assigns subtype labels according to the multi-omics consensus ensemble of muscle-invasive bladder cancer (MIBC) using nearest template prediction. The consensus ensemble identifies 4 integrative consensus subtypes: basal-inflamed, basal-noninflamed, luminal-excluded, and luminal-desert. This package also deploys a 5-immune-gene classifier to refine each basal-like MIBC as either basal-inflamed or basal-noninflamed by a random forest classifier if basal-like classification has already been identified by other approaches (*e*.*g*., CMS, PAM, oneNN, Lund, *etc*.).

## Authors’ contributions

Conceptualization, X. L and J. M; methodology, X. L, L. S, and L. J; formal analysis, X. L, J. M, L. S, J. Z, M. H, W. C, L. X, Y. Z, H. W and L. J; investigation, X. L, J. M, J. Z, M. H, W. C, L. X, S. Y and H. W; writing the original draft, X. L and J. M; visualization, X. L; funding acquisition, C. L and F. Y, and supervision, C. L and F. Y.

## Acknowledgements

We greatly appreciate the patients and investigators who participated in the corresponding medical project for providing data.

